# Identifying the core bacterial and fungal communities within four agricultural biobeds used for the treatment of pesticide rinsates

**DOI:** 10.1101/298331

**Authors:** Jordyn Bergsveinson, Benjamin J. Perry, Claudia Sheedy, Larry Braul, Sharon Reedyk, Bruce D. Gossen, Christopher K. Yost

## Abstract

Bacterial and fungal communities of four pesticide rinsate treatment biobeds constructed in Alberta and Saskatchewan, Canada were profiled via high throughput DNA sequencing to assess the effect of biobed depth and pesticide application on microbial community composition. Biobeds differed in geographical location and biobed design, and composition of pesticide rinsates (including herbicides, fungicides, and insecticides). All biobeds achieved similar treatment efficacy and supported greater bacterial diversity relative to fungal diversity, yet selected for similar abundant bacterial orders of Actinomycetales, Acidobacteria, Rhizobiales, and Sphingobacteriales and fungal taxonomic groups of Dothideomycetes, Eurotiales, Hypocreales, and Sordariales. Biobeds differed in the presence of unique and differentiated genera and operational taxonomic units. Biobed depth did not uniformly impact the diversity and/or the microbial community structure. Overall, pesticide application increased bacterial diversity, but had limited effect on the more variable fungal diversity, therefore suggesting broader implication for the effect of applied fungicides on biobed fungal communities.

**Highlights:**

- Biobeds support diverse bacterial and fungal communities
- Specific “core” bacterial and fungal taxa are abundant in biobeds of different design and treatment
- Microbial diversity is not directly linked with pesticide type or diversity.

## 1. Introduction

A significant threat to surface water and groundwater quality in modern agriculture is the improper handling and disposal of pesticides and pesticide waste, in the form of rinsate from farming equipment and pesticide concentrate containers. One proven method for mitigating the release of these hazardous materials into the environment is by remediating the pesticide rinsate through use of a biobed system (Castillo et al., 2008; Vischetti et al., 2004). Biobeds are comprised of an active “bio” matrix (often termed a “biomixture”), which represents a 2:1:1 ratio by volume of lignin substrate (straw or wood), adsorption material (peat or compost), and topsoil. This matrix provides adsorption capacity and facilitates the biodegradation of pesticide residues through the metabolic capacity of the microbial communities within the biomatrix.

It is well established that the activity of various microbial enzymes, including (mono)oxygenases, laccases, peroxidases, lipases, cellulases, and proteases (Karigar and Rao, 2011) produced by lignin-degrading fungi and bacteria have the capacity to degrade xenobiotics, including pesticides (Castillo et al., 1997, 2000; Kotterman et al., 1998; Tortella et al., 2013; Varsha et al., 2011). The presence and metabolic activity of such microorganisms are known to exist and function within biobed systems (Castillo et al., 2008), and characterization of associated bacterial or fungal communities have been investigated with the aid of denaturing gradient gel electrophoresis (DGGE), coupled with pyrosequencing of the 16S rRNA gene (Holmsgaard, et al., 2017; Marinozzi et al., 2013; Paul et al., 2006). As an improvement to these methods, high throughput DNA sequencing of bacterial 16S rRNA gene and fungal ITS region, provides an opportunity to profile biobed microbial communities at substantially higher resolution, and thus gain greater insight into community stability and diversity, and ultimately aid in future biobed design and optimization.

Agriculture and Agri-Food Canada (AAFC) has led an initiative to establish biobed pesticide treatment systems in Canada. Multiple biobeds have been constructed and are operational within the Canadian prairies. This study focuses on investigating the microbial communities within four specific biobeds located in Alberta and Saskatchewan, Canada. By analyzing multiple biomixture samples collected at varying depths from these biobeds, it is possible to provide increased resolution of potential shifts in the community profile and insight into potential treatment capacity at different depths. While each of the four biobeds received different pesticide residues and are independent environments and/or microbial ecosystems, they do share specific design attributes and thus allow for observations regarding general trends of microbial community composition and stability.

## 2. Materials and Methods

### 2.1. Biobed construction and treatment

The design of each biobed included either one or two “cells”, with pesticide rinsate influent entering and exiting a single biobed (one cell) or flowing through two biobeds in sequence (two cells). One-cell biobeds using straw biomix were constructed at Grande Prairie, AB (below ground; 44 m^2^ surface area) and Vegreville, AB (above ground; 8 m^2^ surface area) (**Table 1**). Two-cell biobeds using wood biomix were constructed in Simpson, SK (above ground; each 4.5 m^2^ surface area) and Outlook, SK (above ground; each 6 m^2^ surface area) (**Table 1**). The biobeds were 1 m deep except at Outlook (OL), which was 0.5 m deep. The effluent from the Grande Prairie (GP) biobed was recirculated through the bed. The GP biobed had a rainfall cover installed early in the operation cycle, which reduced the rinsate volume discharged to the environment by 85%. The biobeds at Vegreville (VG), Simpson (SP) and OL used a low-through design in which effluent not recirculated.

**Table 1.** Overview of biobed design and samples collected for analyses.

| Location | Bed Treatment |  |  | Number of Samples Collected |  |  |  |
| --- | --- | --- | --- | --- | --- | --- | --- |
|  | Year Built; Year Operational | Lignin Source | Design | “Upper”<br>0 – 15 cm | “Lower”<br>15 – 45 cm | “Deep”<br>45 – 100 cm | Total |
| **Grande Prairie, AB**<br>**(“GP”)** | 2012; 2014 | Straw | 1 bed | 8 | 8 | 4 | 20 |
| **Vegreville, AB**<br>**(“VG”)** | 2014; 2015 | Straw | 1 bed | 8 | 8 | 4 | 20 |
| **Outlook, SK**<br>**(“OL”)** | 2012; 2012 | Wood | 2 beds | 8 | 8 | 0<sup>2</sup> | 32<sup>1</sup> |
| **Simpson, SK**<br>**(“SP”)** | 2014; 2015 | Wood | 2 beds | 8 | 8 | 4 | 40<sup>1</sup> |

<sup>1</sup> Collected from Biobed 1 and 2.
<sup>2</sup> Outlook biobed built to shallow depth (0.5 m).

### 2.2. Biobed sampling

Pesticide rinsate biobeds influent and effluent samples were collected throughout the 2015 operational season (**Table A.1**) in 1L amber glass bottles and immediately refrigerated. Samples were shipped to the analytical laboratory in coolers with ice packs and stored at 4°C upon reception until processed.

At the end of the treatment season in 2015 (September), eight composite samples of the biomixture were collected from the Upper (0 – 15 cm deep) and Middle (15 – 45 cm deep) portions of each biobed (**Table 1**) for microbial and pesticide analysis using a soil corer. Also, four samples were collected from the Deep level (45 – 100 cm) in every biobed except OL (**Table 1**). During collection in the field, samples were kept on ice until transport to the laboratory for storage at - 80°C. One soil sample from each biobed prior to pesticide application were similarly collected and processed.

### 2.3. Pesticide analysis

Influent and effluent samples were analyzed for an analytical suite of 142 pesticides. All samples were filtered through glass wool, acidified with concentrated sulfuric acid to pH 2 and extracted by liquid-liquid partitioning with dichloromethane. Extracts were dried with acidified Na_2_SO_4_, concentrated, methylated using diazomethane, transferred to hexane and adjusted to a 10 mL final volume. Esterified extracts were analyzed (2 µL injections) using an Agilent 7890B gas chromatograph with a 7000C QQQ mass selective detector (MSD) in MRM (multiple reaction monitoring). The column was HP-5MS UI 30m × .25mm × .25um, p/n 19091S-433UI. Temperature programming was: 70°C for 2 min, ramp of 25°C/min to 150°C then ramp 3°C/min to 200°C, then ramp of 8°C/min to 280°C for 7 min, with a total analysis time of 38.867 min. One target ion and at least two qualifier ions were monitored, and method blanks were run with each set of samples analyzed. The limit of detection was 0.025 µg for most pesticides, and any detections below these limits were assigned values of ½ LOQ. Limits of quantification and recoveries for all pesticides are included in **Table A.1**.

### 2.4. Environmental DNA (eDNA) isolation and PCR amplicon sequencing

The MoBio Powersoil DNA Isolation kit (MoBio, Carlsbad, CA) was used to extract eDNA from 0.25 g of homogenized biomix material (soil, peat, and wood or straw) from each biobed sample according to the manufacturer’s instructions. Eight DNA samples were prepared from each Upper and Middle sample, and four DNA samples from each Deep sample (Table 1; not available for OL). Sequencing libraries were prepared according to Kozich *et al.* (2013) using PCR primers designed for the v4 hypervariable region of the bacterial 16S rRNA gene (Wu et al., 2015) and PCR primers for the fungal ITS2 region (Schoch et al., 2012) as described by Gweon et al. (2015). DNA sequencing of the PCR amplicons was performed on an Illumina MiSeq platform for both bacterial (v2; 250 bp paired-end) and fungal (v3; 300 bp paired-end) libraries.

### 2.5. Sequence processing and statistical analyses

16S rDNA amplicon sequencing data were processed using Mothur (v.1.39.5; Kozich *et al*. 2013) to remove low-quality reads and chimeric sequences and to perform operational taxonomic unit (OTU) assignment at the 97% identity level, using the SILVA reference bacterial database (v.123; Quast *et al*. 2013) for taxonomic identification. ITS2 amplicon sequencing data were processed using the PIPITS fungal taxonomic assignment software (v.1.4.2; Gweon *et al*. 2015) with the UNITE fungal ITS reference database (v.7.0; Kõljalg *et al.* 2013). Bacterial and fungal taxonomic abundance data for all biobeds were loaded into Calypso web-server software (v.8.46; Zakrzewski et al., 2016) for normalization and visualization. Data were transformed or normalized using cumulative-sum scaling (CSS) with log transformation, discarding samples with less than 1000 sequence reads and excluding taxa with less than 0.01% relative abundance across all samples. Within Calypso, taxonomic abundance data was analyzed for Simpson’s reciprocal diversity metric, similar community composition using non-metric dimensional scaling (NMDS), and for taxa uniquely associated with and/or differentially abundant in specific biobeds using the linear discriminant analysis effect size (LEfSe) method (Segata et al., 2011).

## 3. Results

### 3.1. Pesticide treatment performance

While each biobed experienced individual environmental conditions, only GP was subject to fluctuating operational treatments, given irrigation of this bed with pesticide rinsate was controlled manually and therefore, pesticide application was not consistent (**Table A.1**). Each biobed received pesticide rinsates that differed in the number, concentration, and composition of pesticides (**Fig. 1; Table A.1)**. Common or abundant pesticides across biobeds included auxinic herbicides such as 2,4-D and MCPA (phenoxy), dicamba (benzoic acids), picloram, clopyralid (pyridine acids), fluroxypyr (a synthetic auxin) and triazole fungicides including difenaconazole I and propicanozole. One-cell biobed GP, and two-cell biobed SP received few, or no fungicides in their respective applied rinsates (**Table A.1**). A few recalcitrant pesticides persisted within the biobeds (notably in GP and VG; **Fig. 1**), including mecoprop and clopyralid, a pyridine acid herbicide that is known to persist in dead plant tissue and compost (Haskell, 2003).

**Fig. 1.**
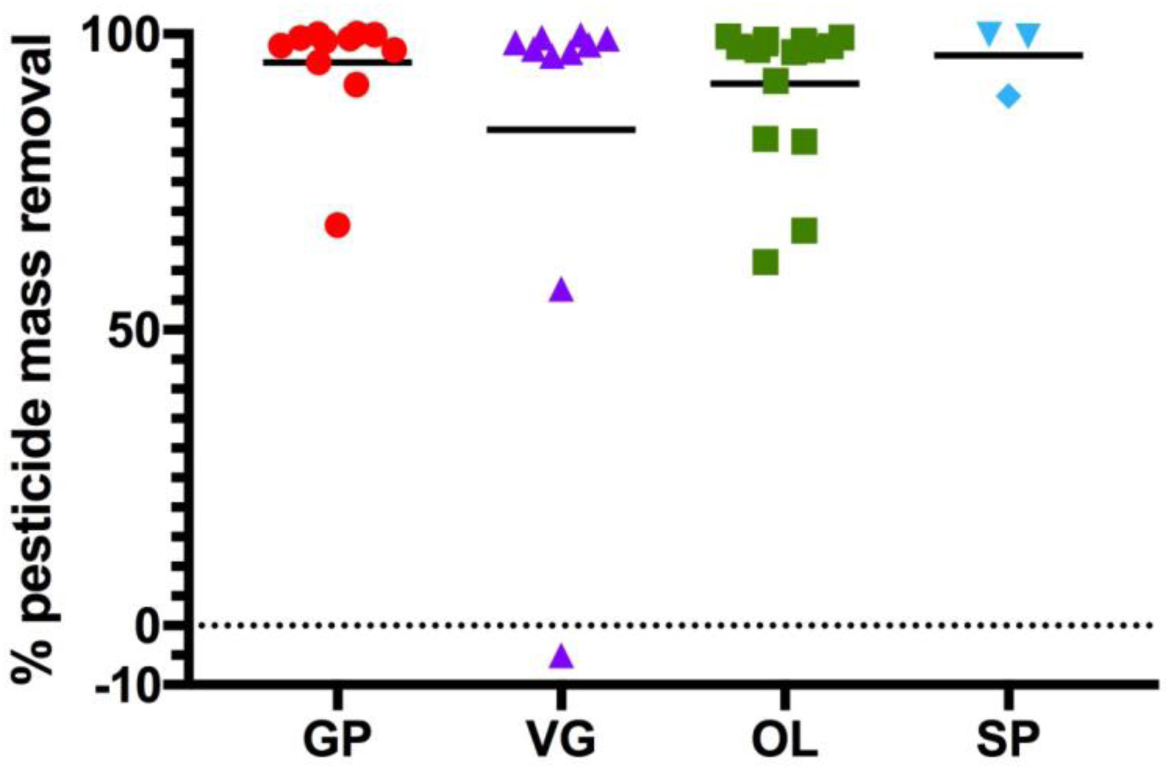
Percent mass removal for individual pesticides present in each biobed. Pesticides present above 0.1 ppb in each biobed are plotted (Table A.1), with average % mass pesticide removal of each bed indicated (horizontal bar).

One-cell biobed systems of GP and VG, both which included straw in their biomix, received similar number of pesticides, with each having 10 pesticides detected above 0.1 ppb (**Fig. 1**; **Table A.1**), however with the important difference that the GP biobed was operated manually, received re-circulated effluent and was covered with a transparent roof, leading to extensive evapotranspiration and highly concentrated effluent. Nonetheless, both biobeds achieved similar overall treatment efficacy rates (total pesticide mass removal rate between 97– 98%; **Table A.1**). While the two-cell biobeds systems of OL and SP both contained woodchips in their biomix and achieved similar overall treatment efficacy rates (99%), they differed in the number and concentration of the pesticides they received, with 39 and 3 unique pesticides detected above 0.1 ppb, respectively. OL and SP biobeds also differed in the % pesticide mass removal performed by their respective first and second biobed cells (**Table A.1**), though ultimately, the presence of BB2 contributed to the removal of pesticide mass and the high overall pesticide mass removal of the systems (99%; **Table A.1**). Despite the recalcitrant nature of few of pesticide compounds, the average percent mass pesticide removal for each biobed is high, regardless of the number and combination of pesticides received (**Fig. 1**)

### 3.2. Bacterial and fungal communities

#### 3.2.1. Diversity

Quality-processed amplicon sequencing reads from each biobed were rarefied independently, resulting in a range of 3,384 to 4,617 bacterial sequences, and 2,816 to 30,274 fungal sequences. Rarefaction curves for each sample reached a plateau, indicating that sequencing efforts captured the complete microbial richness. Examination of microbial community diversity throughout all biobeds was performed using Simpson’s reciprocal metric, which measures both species richness and evenness (**Fig. 2**).

**Fig. 2.**
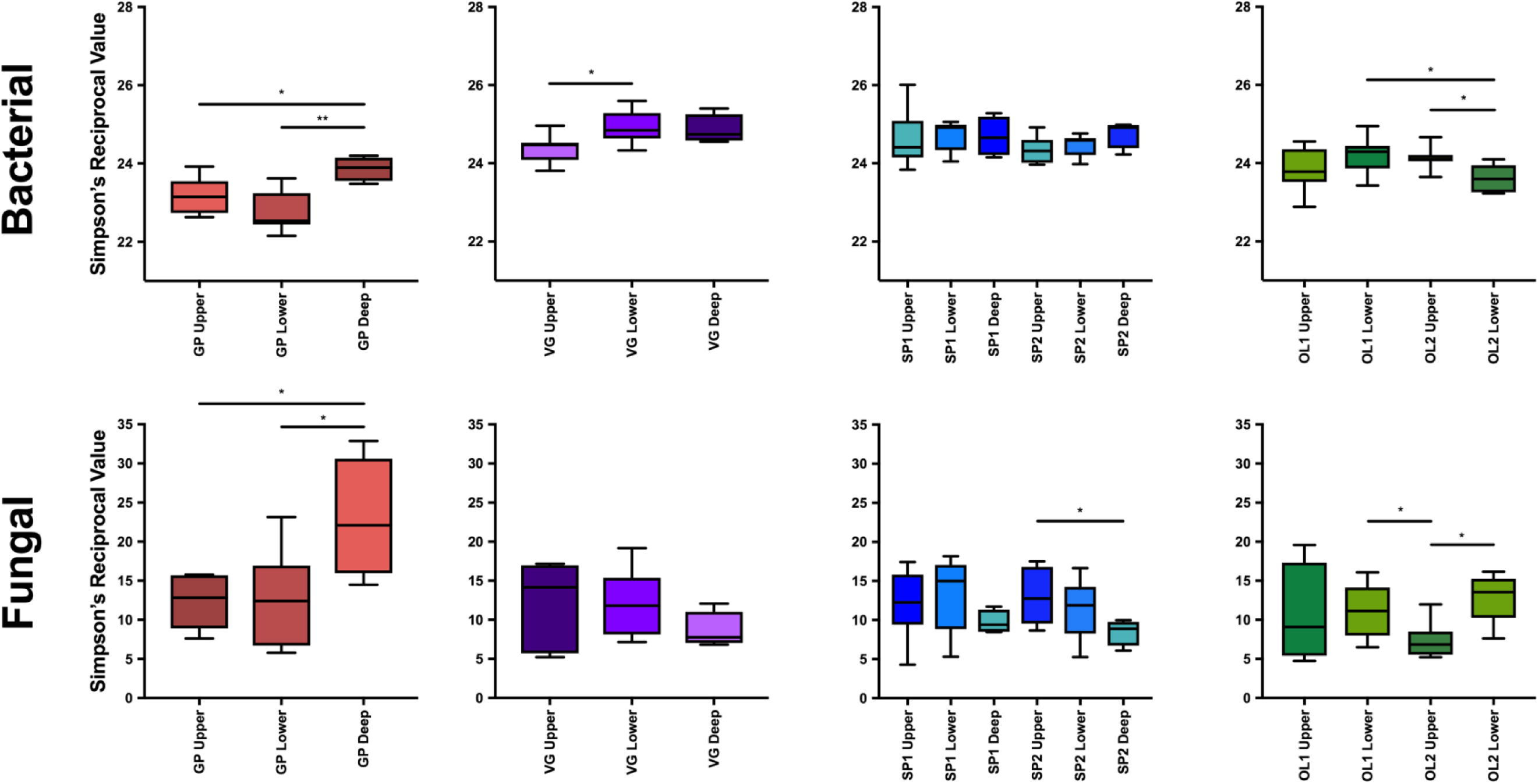
Bacterial and fungal OTU diversity of all collected samples as measured by Simpson’s Reciprocal metric. OTU diversity measures for each sample collected are plotted (upper and lower n= 8; deep = 4). Biobeds 1 and 2 of two-cells systems are referred to as SP1/OL1 and SP2/OL2, respectively. Significant difference between individual sample types within a biobed are indicated within plots (*p* < 0.05).

Each biobed was treated as an independent ecosystem, so direct numerical comparison of diversity measurements should not be over-interpreted. However bacterial communities were generally more diverse than the fungal communities across all four biobeds at every depth (**Fig. 2**). The biobeds had generally similar numbers of bacterial genera across all depths (GP = 456; VG = 447; OL = 397; SP = 477), with inverse Simpson’s values ranging between 23 – 26. Similarly, the biobeds had similar numbers of fungal genera across all depths (GP = 249; VG = 326; OL = 304; SP = 263), but the range in the inverse Simpson’s metric was wider, from 5 – 30 (**Fig. 2**). Also, variance in fungal community diversity was larger amongst samples collected from one biobed location, than for bacterial diversity, a pattern also detected in soil microbial communities (Girvan et al., 2004). Diversity scores and examination of abundant orders (Table A.2) indicated that, relative to bacterial community composition, fungal communities had fewer constituent members (reduced richness), with a subset of these members dominating the general population abundance (reduced evenness) relative to bacterial communities. There was no clear relationship or pattern between diversity of either bacterial or fungal communities and biobed depth (**Fig. 2**).

Bacterial communities of biomixture samples collected from each biobed prior to pesticide application demonstrated markedly lower diversity (16 – 18 Simpson’s Reciprocal Value; data not shown), whereas pre-pesticide application fungal communities exhibited diversity values of 17 – 19, which fall within the range of diversity values exhibited by samples collected at the end of the treatment season.

#### 3.2.2. Community similarity

To assess the uniqueness of biobed communities, non-metric dimensional scaling (NMDS) was used to ordinate the samples from each biobed. NDMS uses a dissimilarity matrix to group similar samples close together. This analysis demonstrated that a similar “core” of classes (or broader taxonomic groups) of both bacteria and fungi were present at high frequency in all four biobeds. However, the genera and OTU (operational taxonomic unit, which functions as an approximation for species) profile of individual biobeds were distinct; as both bacterial and fungal samples from cluster tightly together according to biobed source when genera and OTUs taxa are considered (**Fig. 3**). This differentiation of abundant specific taxa in individual biobeds likely results from the unique pesticide compositions in the influent and differences in the operation and environment of each biobed. For example, samples from the GP biobed exhibit clear differentiation and patterning relative to the other biobeds across all ranks of bacterial and fungal taxa.

**Fig. 3.**
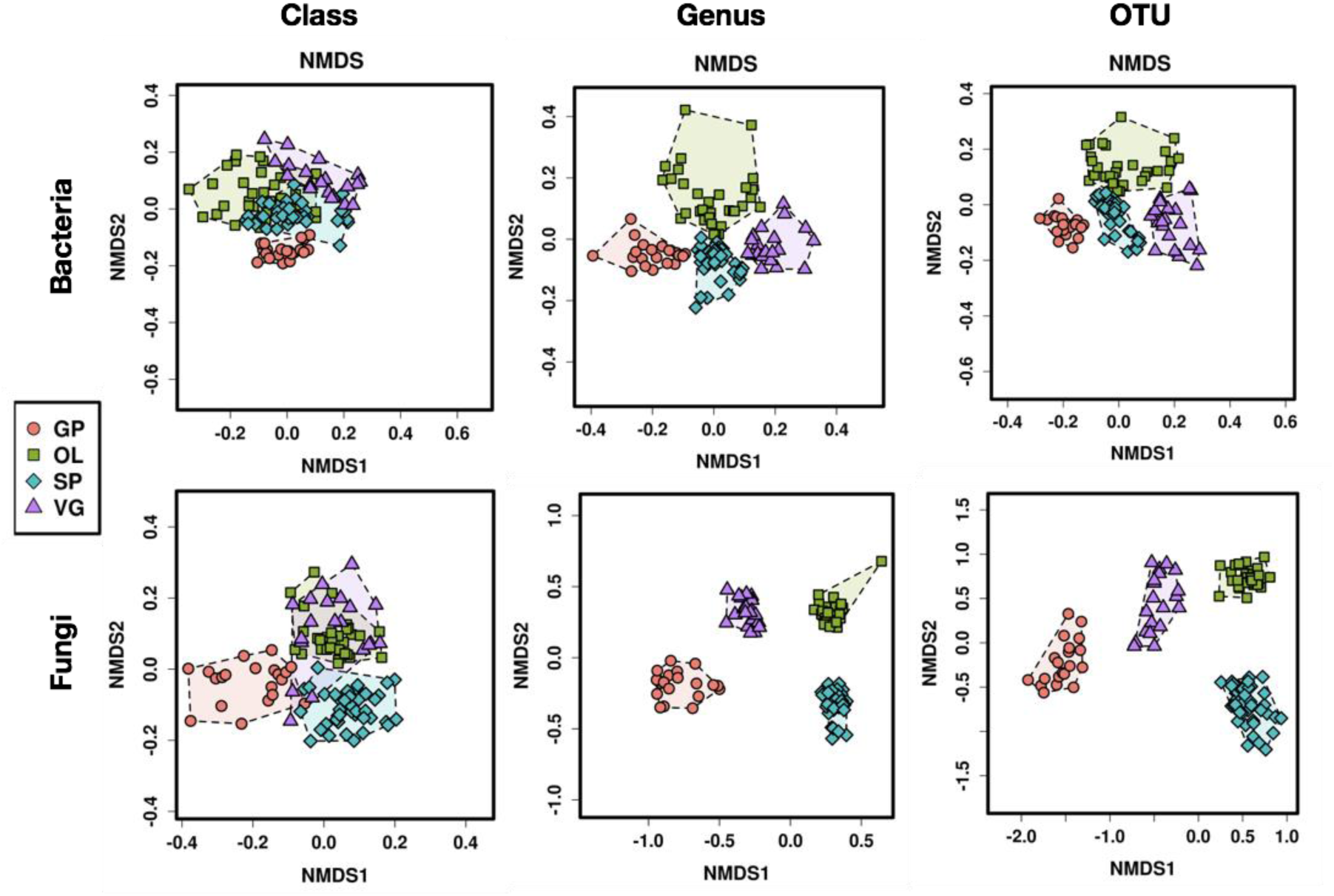
Nonmetric multidimensional scaling (NMDS) analysis of bacterial and fungal biobed communities at level of class, genus, and OTU. Each point in the ordination space is a sample (all of “upper”, “lower”, and “deep” are plotted), and samples that are more similar ordinate closer together. Stress values for all six plots range between <0.05 – 0.09, indicating good representation. Axes and orientation of the plots are arbitrary.

#### 3.3.3. Community profile

Of the top 100 most abundant orders, 60 are shared by all biobeds (**Fig. 4**). Abundant shared bacterial orders include Actinomycetales, Acidobacteria, Rhizobiales, and Sphingobacteriales. Similarly, of the 25 identifiable fungal orders, 17 are shared by all of the biobeds, and include Dothideomycetes, Eurotiales, Hypocreales, and Sordariales (**Table A.2**). The observation that differentiation of microbial biobed communities occurs at lower taxonomic orders indicated that the biobed environment is conducive for growth and development of a cohort of bacterial and fungal orders. To quantify the magnitude of differences at the genus level between biobeds, the top 100 most abundant taxa across biobeds were subjected to LEfSe analysis. In total, 81 bacterial and 50 fungal genera were differentially abundant, based a logarithmic linear discriminant analysis (LDA) score of 2.5 (**Fig. 5**). Notably, fungal genera that were differentially abundant across biobeds exhibited larger LDA scores (range 3.5 – 4.8 log LDA) relative to the differentially abundant bacterial genera (range 2.8 – 3.5 log LDA). The most differentially abundant genera included in *Acidobacteria* Gp1 and *Rhizomicrobium* in VG and *Cytophaga* and *Ohtawekwangia* in GP. The most differentially abundant fungal genera could not be classified to the genus level, however *Talaromyces* in SP, *Preussia* in GP and *Scutellinia* in OL all exhibited large log LDA scores (**Fig. 5**).

**Fig. 4.**
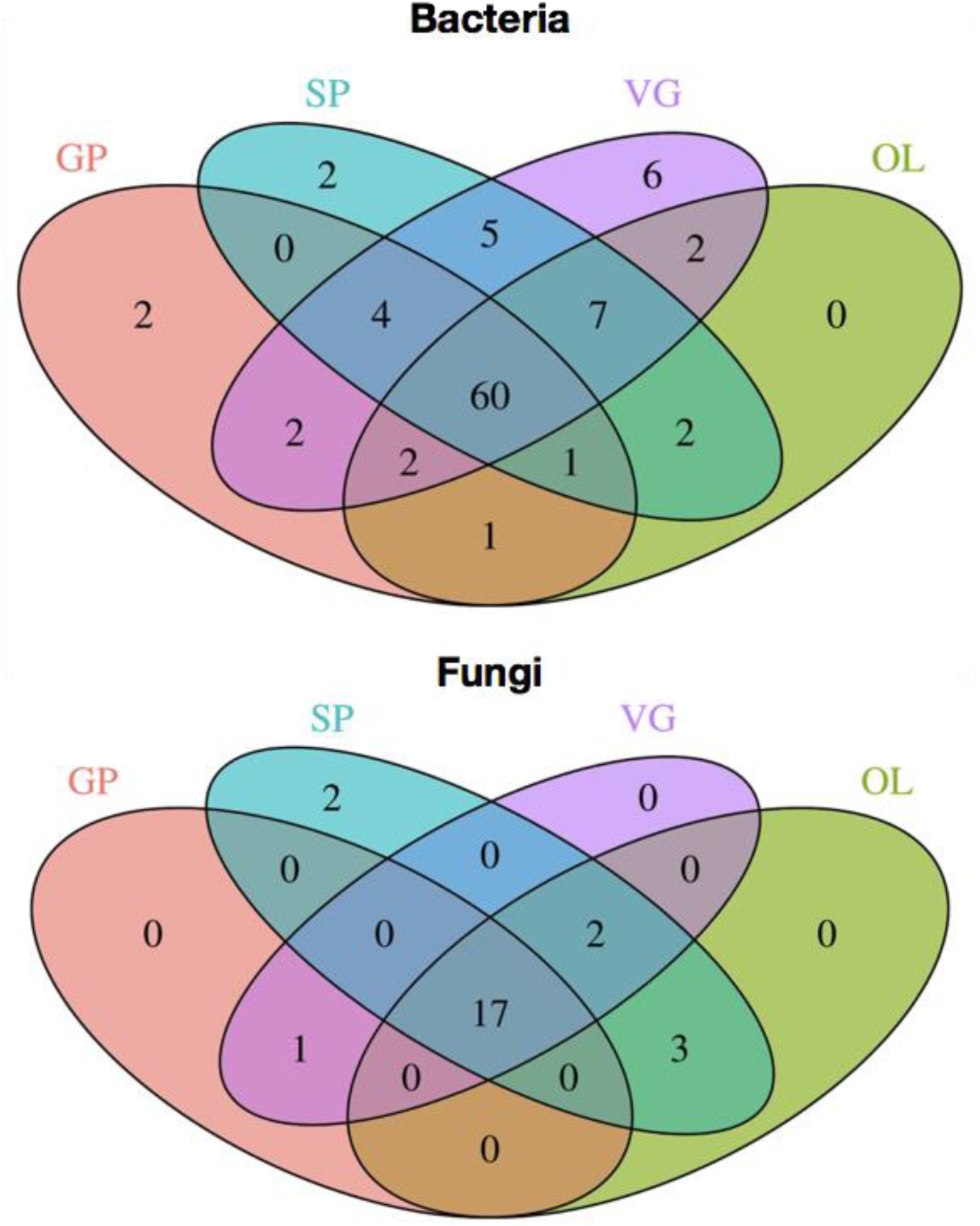
Shared and unique bacterial and fungal orders for each biobed. A total 96 and 25 distinct and identifiable bacterial and fungal orders, respectively, were identified. “Core” taxa are shared by all four biobeds.

**Fig. 5.**
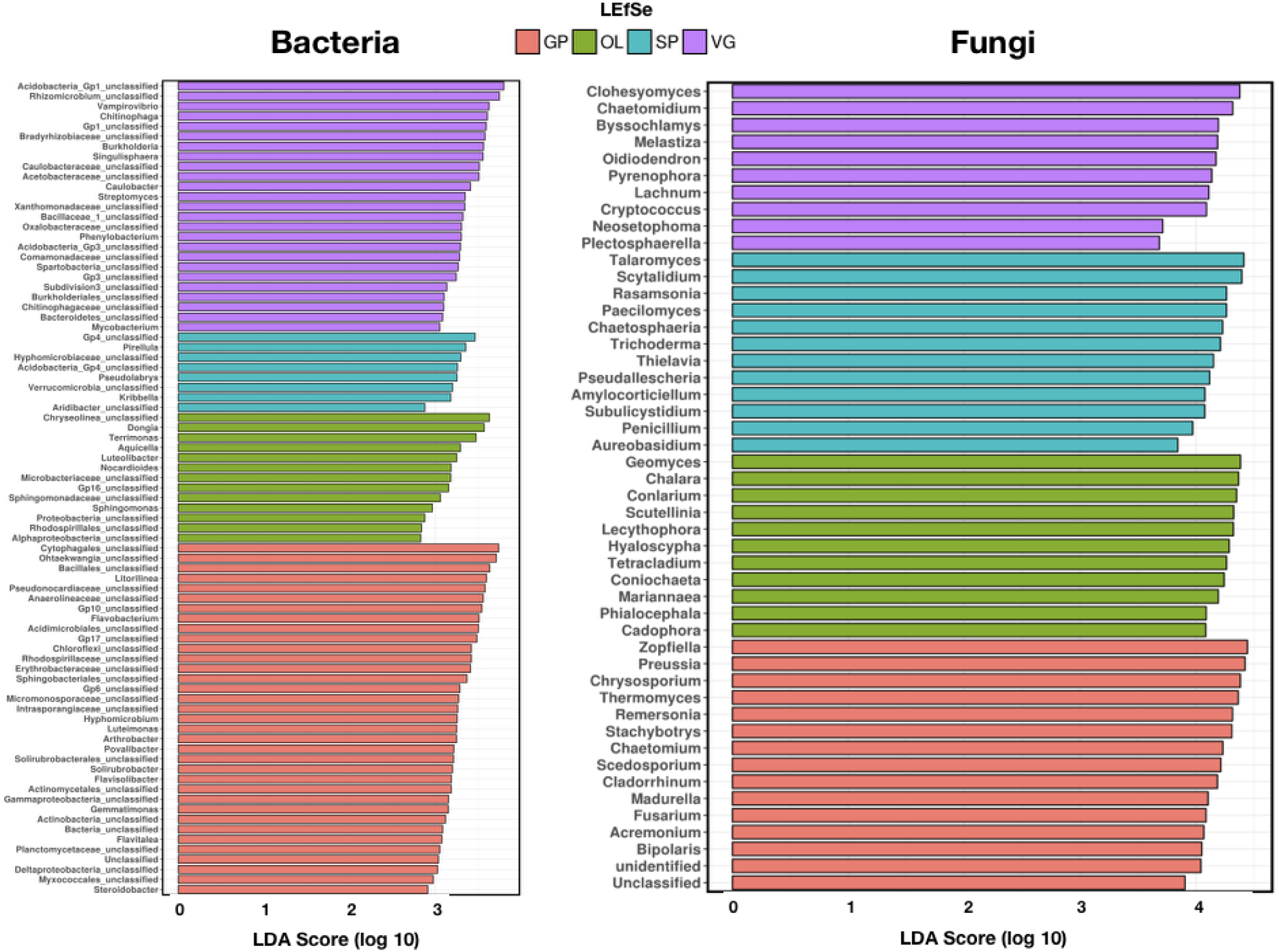
LEfSe discovery of differentially abundant or distinct bacterial and fungal genera between biobeds. For each biobed, listed taxa are significantly (*p* < 0.05) increased in abundance compared to other biobeds, using a logarithmic of LDA score threshold of 2.0. LDA indicates the effect size of each differentially abundant genera for a particular group. The top 100 most abundant taxa were considered for analysis.

## 4. Discussion

Despite the unique operation and design parameters of each biobed, the composition and dynamics of the microbial communities assessed at the end of the operational season were generally similar across sites (**Table 1**). Each biobed supported diverse bacterial and fungal communities, with greater bacterial diversity relative to fungal diversity (**Fig. 2** and **3**). Despite intra-biobed sample variance, and relative differences in diversity, both bacterial and fungal communities in each biobed were similar at higher taxonomic levels (**Fig. 3** and **4**).

Use of high-throughput DNA sequencing allowed for increased resolution of the complexity of both bacterial and fungal communities. Indeed, rarefaction assessments indicated that the entire range of diversity of the microbial communities was captured. This represents an improvement over previous studies of bacterial communities using DGGE and pyrosequencing of 16S rRNA genes, which did not achieve adequate coverage of the community. However, the results of the current study generally support the pattern of bacterial community structure identified using pyrosequencing of 16S rRNA gene fragments (Holmsgaard et al., 2017).

The most dominant bacterial phyla within each biobed were Proteobacteria, followed by Actinobacteria, Bacteroidetes, Chloroflexi, Verrucomicrobia, Firmicutes, and Gemmatimonadetes (Table A.2). The most dominant fungal groups within each biobed were the class Dothideomycetes, and orders of Hypocreales and Sordariales. Fungal communities exhibited greater variation in both diversity and taxa abundance at different depths in the biomixture relative to bacterial communities (**Fig. 2** and **3**).

Bacterial orders frequently associated with the metabolic characteristics that may be important in the biobed environment (Actinomycetales, Acidobacteria Gp6, Bacteroidetes, Rhizobiales and Sphingobacteriales) were among the top 10 most abundant bacterial orders in each biobed (**Fig. 4**; **Table A.2**). For example, Actinomycetales commonly produce monooxygenases and phenoloxidases (De Schrijver and DeMot, 1999), and additionally have a filamentous growth form that helps them to colonize the biomixture effectively and produce long-lived spores under adverse conditions (Briceño et al., 2013). Also, they have less stringent requirements for carbon and nitrogen compared to fungi, which facilitates more metabolic activity independent of the biomixture (Briceño et al., 2013). Other orders include Bacteroidetes, which have been shown to develop on the surface leaves and stems of pesticide-treated plants (Zhang et al., 2009) and Rhizobiales which include a large number of nitrogen-fixing or sulfate-reducing species that have been implicated in the degradation of recalcitrant pesticides (Chaussonnerie et al., 2016).

Identification of the fungal community in each biobed was challenging, due to limitations in fungal identification databases, which make resolution of OTUs beyond the taxonomic rank of phylum difficult (Kõljalg *et al.* 2013). The most abundant fungal organisms in all biobeds (**Table A.2)** were OTUs belonging to the class Dothideomycetes. This is the largest class of fungi, both for number of species and for ecological and biological diversity. Species within this group appear to have a preference for habitats with a high concentration of organic matter, where they participate in biomass conversion (Shrestha et al., 2011). The next most abundant OTUs were in the order Hypocreales, which are distributed worldwide, and show high species diversity and include a broad range of plant associated fungi (Rogerson, 1970; Rossman et al., 1999). Other common OTUs were from the Eurotiales and Sordariales, which are also ubiquitous in soil and/or decaying organic matter, with Eurotiales being known to adapt to extreme environmental conditions (Taylor et al., 2009).

There was no a strong relationship between the number of pesticides applied to a biobed and overall microbial diversity or core composition, based on the community structure and profile of each biobed (**Fig. 2, 3**, and **4**). Notably, the fungal community profiled in GP biobed, which received no fungicide and experienced infrequent rinsate application, was notably more variable at different depths, relative to other biobeds (**Fig. 2**). Further, the application of fungicide in beds VG and OL did not greatly influence the diversity of the fungal community when samples collected pre- and post-rinsate application were compared (data not shown). It is possible that as the samples were collected at the end of the operational season, the presence and effect of applied fungicides had been reduced, therefore returning the fungal community diversity to pre-application range. This is supported by previous work which demonstrated a “threshold” relationship between fungal diversity and concentration of applied fungicide, with low-level concentration having limited impact on observed diversity (Howell et al., 2014). In contrast, it has been reported that bacterial communities do shift in composition and structure over time, especially after pesticides are applied (Holmsgaard et al., 2017). Despite the anticipated shift in bacterial community profile and diversity in response to pesticide application, post-application bacterial diversity and phylogenetic characteristics were similar across the range of biobeds. This indicated that, regardless of differences in the type and concentration of applied pesticides applied, the biobed environment is likely selective for bacteria, and to some extent fungi taxa, with specific physiological or metabolic characteristics, even though there is substantial differentiation at the genus and species levels (**Fig. 3** and **5**).

Genomics technologies have increased our ability to study complex microbial ecosystems. Continued microbial community profiling of biobed systems, with a focus on functional metagenomics, will help to elucidate parameters that affect pesticide removal, and ultimately aid in improving biobed construction and management.

## 5. Conclusions

Biobeds were shown to provide an effective system for management of pesticide rinsate on the Canadian prairies, with efficacy of > 90%. The current study demonstrated that efficient removal of pesticide mass is possible in biobeds of diverse construction and composition. Analysis of microbial communities in biobeds that differed in design, environmental parameters and management protocols demonstrated that a stable cohort of bacterial and fungal taxa, adapted to an environment with high abundance of pesticides, became abundant in each biobed by the end of the operating season, but that these taxa were represented by different OTUs at each site.

## Acknowledgements

This study was funded by an Agriculture and Agri-Food Canada Growing Forward Grant. Pesticide residue analysis was performed at AAFC’s trace residue analysis laboratory in Lethbridge, Alberta. The authors would like to acknowledge contributions to pesticide analysis from D. Nilsson.

## Appendices

**Table A.1**. 2015 biobed pesticide data.

**Table A.2**. Abundance of bacterial and fungal phyla across biobeds.

## References

1. Briceño, G., Pizzul, L., Diez, MC. 2013. Chapter 10: Biodegradation of pesticides by Actinobacteria and their possible application in biobed systems. In: Actinobacteria: Application in bioremediation and production of industrial enzymes. Eds: Amoroso, M.J., Benimeli, C.S., Cuozzo, S.A. CRC Press: Boca Raton, FL. pp. 165–90.

2. Castillo, M.P, Ander, P., Stenström, J. 1997. Lignin and manganese peroxidase activity in extracts from straw solid substrate fermentations. Biotechnol. Tech. 11, 701–6.

3. Castillo, M.P., Ander, P., Stenström J, Torstensson L. 2000. Degradation of the herbicide bentazon as related to enzyme production by *Phanerochaete chrysosporium* in a solid substrate fermentation system. World J. Microbiol. Biotechnol. 16, 289–95.

4. Castillo, M.P., Torstensson, L., Stenström, J. 2008. Biobeds for environmental protection from pesticide use — a review. J. Agric. Food. Chem. 56, 6206–19.

5. Chaussonnerie, S., Saaidi, P.L., Ugarte, E., Barbance, A., Fossey, A., Barbe, V., et al. 2016. Microbial degradation of a recalcitrant pesticide: chlordecone. Front. Microbiol. 7, 2025. https://doi.org/10.3389/fmicb.2016.02025

6. De Schrijver, A., De Mot, R. 1999. Degradation of pesticides by actinomycetes. Crti. Rev. Microbiol. 25, 85–119.

7. Girvan, M.S., Bullimore, J., Ball, A.S., Pretty, J.N., Obsorn, A.M. 2004. Responses of active bacterial and fungal communities in soils under winter wheat to different fertilizer and pesticide regimens. Appl. Environ, Microbiol. 70, 2692–701.

8. Gweon, H.S., Oliver, A., Taylor, J., Booth, T., Gibbs, M., Read, D.S., Griffiths, R.I., Schonrogge, K. 2015. PIPITS: and automated pipeline for analyses of fungal internal transcribed spacer sequences from the Illumina sequencing platform. Meth. Ecol. Evol. 6, 973.

9. Haskell, D.E. 2003. California Department of Pesticide Regulation: Clopyralid in Compost. Proc. Calif. Weed Sci. Soc. 55, 163–6.

10. Holmsgaard, P.N., Dealtry, S., Dunon, V., Heuer, H., Hansen, L.H., Springael, D., Smalla, K., Riber, L., Sørensen, S.J. 2017. Response of the bacterial community in an on-farm biopurification system, to which diverse pesticies are introduced over an agrictulral season. Environ. Pollut. 229, 854–62.

11. Howell, C.C., Hilton, S., Semple, K.T., Bending, G.D. 2014. Resistance and resilience responses of a range of soil eukaryote and bacterial taxa to fungicide application. Chemosphere 112, 194–202.

12. Karigar, C.S., Rao, S.S. 2011. Role of microbial enzymes in the bioremediation of pollutants: A review. Enz. Res. vol. 2011, Article ID 805187, 11 pages, http://dx.doi.org/10.4061/2011/805187

13. Kõljalg, U., Nilsson, R.H., Abarenkov, K., Tedersoo, L., Taylor, A.F.S., Bahram, M. et al. 2013. Towards a unified paradigm for sequence-based identification of fungi. Mol. Ecol, 22, 5271.

14. Kotterman, M.J.J., Vis, E.H., Field, J.A. 1998. Successive mineralization and detoxification of benzo[a]pyrene by the white rot fungus *Bjerkandera* sp. strain BOS55 and indigenous microflora. Appl. Environ. Microbiol. 64, 2853–58.

15. Kozich, J.J., Westcott, S.L., Baxter, N.T., Highlander, S.K., Schloss, P.D. 2013. Development of a dual-index sequencing strategy and curation pipeline for analyzing amplicon sequence data on the Miseq illumina sequencing platform. Appl. Environ. Microbiol. 79, 5112.

16. Marinozzi, M., Coppola, L., Monaci, E., Karpouzas, D.G., Papadopoulou, E., Menkissoglu-Spiroudi, U., Vischetti, C., 2013. The dissipation of three fungicides in a biobed organic substrate and their impact on the structure and activity of the microbial community. Environ. Sci. Pollut. Res. 20, 2546e2555.

17. Paul, D., Pandey, G., Meier, C., van der Meer, J.R., Jain, R.K. 2006. Bacterial community structure of a pesticide-contaminated site and assessment of changes induced in community structure during bioremediation. FEMS Ecol. 57, 116–27.

18. Quast, C., Pruesse, E., Yilmaz, P., Gerken, J., Schweer, T., Yarza, P., Peplies, J., Glöckner, F.O. 2013. The SILVA ribosomal RNA gene database project: improved data processing and web-based tools. Nucl. Acids Res. 41, D590.

19. Rogerson, C.T. 1970. The Hypocrealean fungi (Ascomyceters, Hypocreales). Mycolog. 62, 865–910.

20. Rossman, A.Y., Samuels, G.J., Rogerson, C.T., Lowen, R. 1999. Genera of Bionectriaceae, Hypocreaceae and Nectriaceae (Hypocreales, Ascomycetes). Stud. Mycol. 42: 1–248.

21. Schloss, P.D., Westcott, S.L., Ryabin, T., Hall, J.R., Hartmann, M., Hollister, E.B., Lesniewski, R.A., Oakley, B.B., Parks, D.H., Robinson, C.J., Sahl, J.W., Stres, B., Thallinger, G.G., Van Horn, D.J., Weber, C.F. 2009. Introducing mothur: Open-source, platform-independent, community-supported software for describing and comparing microbial communities. Appl. Environ. Microbiol. 75, 7537–41.

22. Schoch, C.L., Seifert, K.A., Huhndorf, S., Robert, V., Spouge, J.L., Levesque, C.A., Chen, W., Fungal Barcoding Consortium. 2012. Nuclear ribosomal internal transcribed spacer (ITS) region as a universal DNA barcode marker for Fungi. Proc. Nat. Acad. Sci. 109, 6241.

23. Segata, N., Izard, J., Waldron, L., Gevers, D., Miropolsky, L., Garrett, W.S., Huttenhower, C. 2011. Metagenomic biomarker discovery and explanation. Genom. Biol, 12, R60.

24. Shrestha, K., Shrestha, P., Walsh, K.B., Harrower, K.M., Midmore, D.J. 2011. Microbial enhancement of compost extracts based on cattle rumen content compost-characterization of a system. Bioresour. Technol. 102, 8027–34.

25. Taylor, T.N., Taylor, E.L., Krings, M. 2009. 3 – Fungi, Bacteria, and Lichens. In: Paleobotany, 2^nd^ Edition. Academic Press, London, pp. 71–119.

26. Tortella, G.R., Mella-Herrera, R.A., Sousa, D.Z., Rubilar, O., Acuña, J.J., Briceño, G., Diez, M.C. 2013. Atrazine dissipation and its impact on the microbial communities and community level physiological profiles in a microcosm simulating the biomixture of on-farm biopurification system. J. Haz. Mat. 260, 459–67.

27. Varsha, Y.M., Naga, D.C.H., Chenna, S. 2011. An emphasis on xenobiotic degradation in environmental cleanup. J. Bioremed. Biodegrad. S11, 001. doi: 10.4172/2155-6199.S11-001

28. Vischetti, C., Capri, E., Trevisan, M., Casucci, C., Petrucci, P. 2004. Biomassebed: a biological system to reduce pesticide point contamination at farm level. Chemosphere. 55, 823–8.

29. Wu, L., Wen, C., Qin, Y., Yin, H., Tu, Q., Van Nostrand, J.D., Yuan, T., Yuan, M., Deng, Y., Zhou, J. 2015. Phasing amplicon sequencing on Illumina Miseq for robust environmental microbial community analysis. BMC Microbiol, 15, 125.

30. Zakrzewski, M., Proietti, C., Ellis, J., Hasan, S., Brion, M., Berger, B., Krause, L. 2016. Calypso: A user-friendly web-server for mining and visualizing microbiome-environment interactions. Bioinformatics. 33, 782–3.

31. Zhang, B., Bai, Z., Hoefel, D., Tang, L, Wang, X., Li, B., Li, Z., Zhuang, G. 2009. The impacts of cypermethrin pesticide application on the non-target microbial community of the pepper plant phyllosphere. Sci. Tot. Environ. 407, 1915–22.

